# Yeast transformation efficiency is enhanced by TORC1- and eisosome-dependent signalling

**DOI:** 10.1101/244889

**Authors:** Sheng-Chun Yu, Florian Kuemmel, Maria-Nefeli Skoufou-Papoutsaki, Pietro D Spanu

**Affiliations:** Department of Life Sciences, South Kensington Campus, Imperial College London, London, SW7 2AZ, United Kingdom.

## Abstract

Transformation of baker’s yeast (*Saccharomyces cerevisiae*) plays a key role in several experimental techniques, yet the molecular mechanisms underpinning transformation are still unclear. Addition of amino acids to the growth and transformation medium increases transformation efficiency. Here, we show that target of rapamycin complex 1 (TORC1) activated by amino acids enhances transformation via ubiquitin-mediated endocytosis. We created mutants of the TORC1 pathway, α-arrestins, and eisosome-related genes. Our results demonstrate that the TORC1-Npr1-Art1/Rsp5 pathway regulates yeast transformation. Based on our previous study, activation of this pathway results in a 13-fold increase in transformation efficiency, or greater. Additionally, we suggest DNA is taken up by domains at the membrane compartment of Can1 (MCC) in the plasma membrane formed by eisosomes. Yeast studies on transformation could be used as a platform to understand the mechanism of DNA uptake in mammalian systems, which is clinically relevant to optimise gene therapy.

## INTRODUCTION

Yeast transformation is the process by which exogenous DNA is introduced into the cell. It is a powerful tool of molecular biology research, for example in the yeast two-hybrid (Y2H) system for detection of protein-protein interactions (Fields and Song 1989). Highly efficient protocols for chemical transformation have been established (Gietz 2015) but the molecular mechanisms underlying yeast transformation are not well understood. Several studies over the last four decades have investigated how DNA passes through the cell wall, through the plasma membrane (PM), and subsequently reaches the nucleus (Kawai, Hashimoto et al. 2010, Mitrikeski 2013). Foreign DNA is most likely to be engulfed via endocytic membrane invagination and several mutants involved in endocytosis show low transformation efficiencies (Kawai, Pham et al. 2004).

Ubiquitination of plasma membrane proteins can serve as an internalisation signal for endocytosis (Toret and Drubin 2006). In this way, the cell can downregulate receptors or transporters via transport to endosomes and lysosomal degradation (Ghaddar, Merhi et al. 2014). In yeast, this process is mediated by the Rsp5 ubiquitin ligase that requires adaptor proteins for recruitment to the specific plasma membrane targets. Proteins that bind to Rsp5 and promote this function, include the arrestin-related trafficking adaptors (ARTs) such as Art1, Art3 and Bul1 (Yashiroda, Oguchi et al. 1996, Lin, MacGurn et al.). Several amino acid transporters are internalised by this form of endocytosis, including the general amino acid permease (Gap1) and the arginine-specific permease (Can1). The latter are targets of Bul1 and Art1, respectively (Ghaddar, Merhi et al. 2014). Phosphorylation of Art1 and Bul1/2 by the Npr1 kinase cause translocation of Art1 from the plasma membrane to the Golgi apparatus and binding of Bul1 to the inhibitory 14-3-3 proteins. This prevents internalisation of the plasma membrane permeases (MacGurn, Hsu et al. 2011, O’Donnell 2012).

The target of rapamycin complex 1 (TORC1) is highly conserved among eukaryotes and functions as a master regulator of cell growth and metabolism through its own as well as downstream protein kinases. TORC1 activity depends on nutrient availability, and amino acids are potent stimulators *in vivo* (Conrad, Schothorst et al. 2014, Dokudovskaya and Rout 2015, González and Hall 2017). When active, TORC1 promotes ubiquitin-mediated endocytosis by inhibiting Npr1, which in turn is a negative regulator of α-arrestins acting in endocytic cargo sorting. Npr1 phosphorylates Art1 and this prevents Art1 from associating with the PM. In this way, the target of rapamycin (TOR) pathway connects amino acid sensing with endocytosis (MacGurn, Hsu et al. 2011).

The downstream plasma membrane targets of Art1 include nutrient permeases that accumulate in the membrane compartment of Can1 (MCC). The membrane composition of MCC differs from other membrane compartments by its higher content of ergosterol (Grossmann, Opekarová et al. 2007). Eisosomes, first discovered in 1963 in early electron microscopy studies (Moor and Muhlethaler 1963), are cytosolic multi-protein complexes that form 50–300 nm deep furrow-like invaginations of the plasma membrane associated with the MCC region (Stradalova, Stahlschmidt et al. 2009). Eisosomes act as a hub for various signalling pathways and they may play a role in endocytosis, although their exact function is not well understood (Walther, Brickner et al. 2006, Fröhlich, Moreira et al. 2009, Douglas and Konopka 2014).

Recently, we developed a yeast transformation protocol called SuccessAA (Yu, Dawson et al. 2016). Using this method, we found that adding nutrients to the transformation and competence reagents substantially increased transformation efficiencies. We speculated that the mechanism underlying this effect was due to the activation of the TORC1 complex, which in turn promotes DNA uptake via ubiquitin-mediated endocytosis.

The aim of our study was to investigate the molecular mechanisms of yeast transformation and the events that lead to the increase in transformation efficiency by addition of nutrients. We found that mutations of endocytic components resulted in changes in transformation efficiencies supporting the hypothesis that TORC1 and ubiquitin-mediated endocytosis is key to yeast transformation. Moreover, the boosting effect was observed in several distinct strains, highlighting the potential for the general application of the SuccessAA protocol to budding yeast transformation.

## RESULTS

### The addition of amino acids results in increased transformation efficiency in different S. *cerevisiae* strains

The applicability of the SuccessAA protocol to different budding yeast strains was examined by transforming four S. *cerevisiae* strains with a 13.8 kb plasmid (Figure 1A) namely THY.AP4, BY4743, W303–1A and MaV203. In all cases, the transformation efficiency after the addition of nutrients was substantially higher than without nutrient addition. The observed increases in efficiency were 15-fold (p=0.0079); 6-fold (p=0.0079) and 37-fold (p=0.0079) for THY.AP4, BY4743 and W303–1A, respectively. We deduce that the genetic requirements for this effect are likely to be conserved in S. *cerevisiae*. Therefore, adding nutrients during yeast transformation may provide a generally applicable method to boost transformation efficiency for budding yeast.

**Figure 1.**
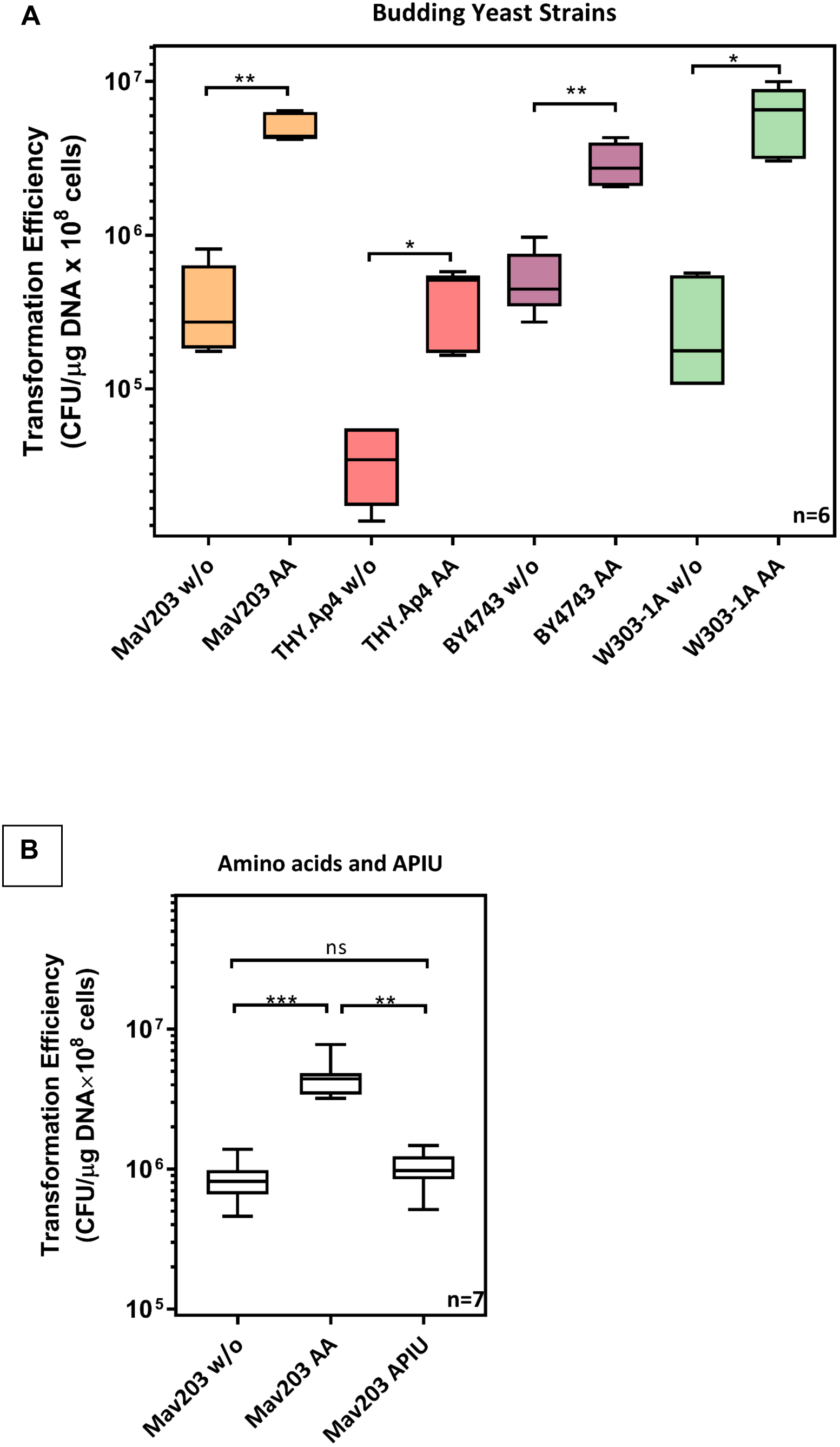
Amino acid addition results in higher transformation efficiency in different budding yeast strains. (A) Four budding yeast strains, namely MaV203, THY.Ap4, BY4743, and W303–1A, were used to test the effect of nutrient addition on boosting yeast transformation efficiency. Each one of the strains was transformed with or without nutrient supplement (drop-out leucine) during the transformation process, followed by three-day culture on 1x YPAD drop-out leucine plates and then the corresponding transformation efficiencies were calculated. Statistical significance was assessed by using the Mann-Whitney test. Nutrient addition caused a significant increase in transformation efficiency in all four yeast strains. Results are from six independent biological replicates. (B) The MaV203 yeast strain was used to test if the boost effect was caused by either amino acids or by four other nutrients, namely, adenine, p-aminobenzoic acid, inositol, and uracil (APIU). MaV203 was transformed without nutrient addition, with amino acid addition (drop-out leucine), or with APIU addition, followed by three-day culturing on 1x YPAD drop-out leucine plates. The corresponding efficiencies were then assessed by the Kruskal-Wallis test, followed by Dunn’s post-hoc test. Amino acid addition resulted in significantly higher transformation efficiencies. Results are from seven independent biological replicates.

To determine which of the nutrients added in the SuccessAA protocol are necessary for the boosting of competence, we tested the effect of a mixture of all amino acids (AA) or Adenine, p-aminobenzoic acid, Inositol, and Uracil (APIU). The addition of only APIU did not cause any significant change in the transformation efficiency (Figure 1B) while amino acids caused the same increases seen in previous experiments where a complex nutrient supplement was employed.

### TORC1-regulated endocytosis is required for enhanced transformation efficiency

Previous studies demonstrated that amino acids in the growth medium activate the TORC1 complex which in turn regulates ubiquitin-mediated endocytosis of nutrient permeases via Npr1 and Art1 (MacGurn, Hsu et al. 2011). Here, we tested the hypothesis that altering the rate of endocytosis by addition of amino acids leads to increased DNA uptake. To achieve this, we created mutant strains lacking components of the TORC1-dependent endocytic pathway (Figure 2), and analysed the effect this had on transformation efficiency (Figure 3). We found that transformation was no longer influenced by addition of amino acids to the medium and transformation mix when either *tco89* (the core subunit of TORC1, p=0.9619) or *art1* (p= 0.9983) had been deleted. Conversely, the efficiency increases when *npr1*, a negative regulator of TORC1 mediated endocytosis, was missing. Note, this effect was visible even in the absence of any additions to the medium (Tukey’s multiple comparisons test, p=0.0021 (wild type yeast without amino acid addition vs *npr1Δ* without amino acid addition), but it was further enhanced when amino acids were supplied (p<0.0001 (wild type yeast with amino acid addition vs *Δnpr1* with amino acid addition).

It has previously been reported that when TORC1 is inactivated, Npr1 stabilizes the yeast plasma membrane general amino acid permease Gap1, by phosphorylating alpha arrestin-like adaptors (Bul1/2); this leads to binding of 14–3–3 proteins and cellular re-localization, which antagonises ubiquitin-mediated endocytosis (Merhi and Andre 2012). Phosphorylation of α-arrestins or arrestin-like adaptors (Art1, Art2, Art3, Art5, Art6, and Bul1/2) increases in rapamycin-treated yeast cells (Iesmantavicius, Weinert et al. 2014). Art2 is an arrestin that does not regulate amino acid induced endocytosis (Nikko and Pelham 2009) and Art5 targets a permease for inositol, which is not involved in the transformation enhancing effect we observed. Furthermore, the activity and the phosphorylation of Art3, but not Art6, are directly regulated by Npr1 (O’Donnell 2012).

**Figure 2.**
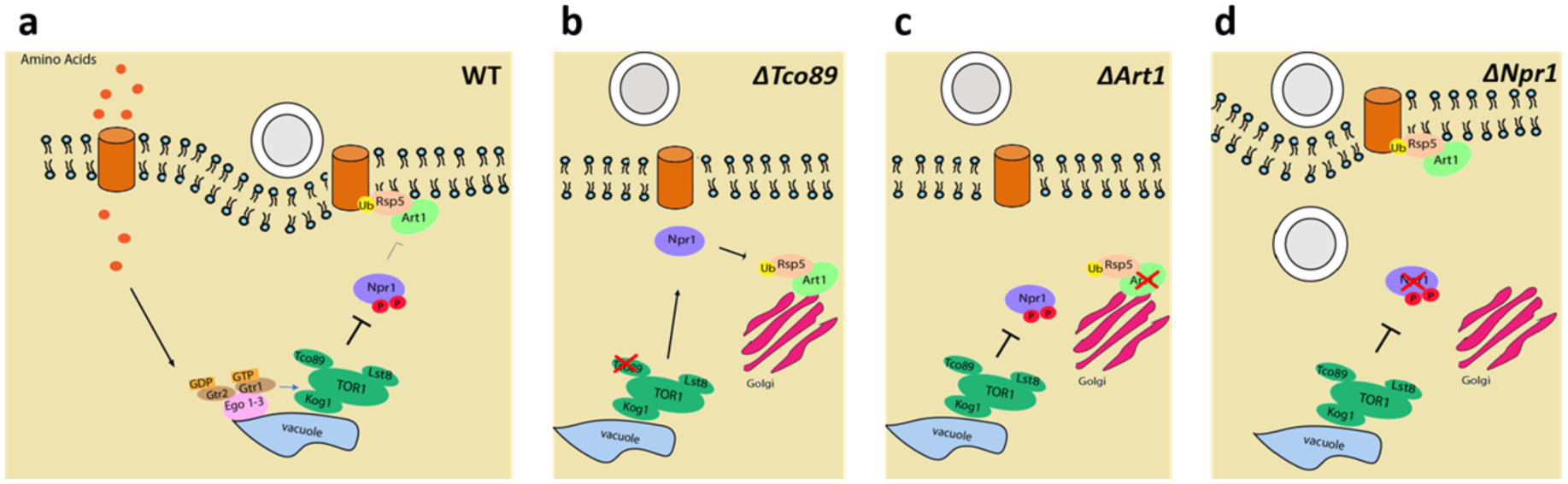
Illustration of the signalling pathways connecting perception of amino acids and endocytosis in wild-type, *Δtco89, Δart1* and *Δnpr1* cells. **(A)** In wild type yeast, intracellular amino acids stimulate TORC1, via the Ego 1–3 complex. TORC1 in turn inhibits Npr1 by phosphorylation, allowing the alpha arrestin adaptor Art1 and ubiquitin ligase Rsp5 to be recruited in the plasma membrane and bind to amino acid transporters, leading to their ubiquitination, subsequent endocytosis and simultaneous plasmid DNA uptake. **(B)** When *tco89* is deleted, TORC1 signalling is impaired, the Npr1 kinase is active and phosphorylates Art1 leading to Art1/Rsp5 translocating to the Golgi-apparatus. Amino acid permeases endocytosis and simultaneous plasmid DNA uptake are hindered, as the amino acid permeases will not be ubiquitinated. **(C)** When *art1* is deleted, Rsp5 cannot be recruited at the plasma membrane to bind to and ubiquitinate amino acid permeases Therefore, endocytosis of nutrient permeases and simultaneous plasmid DNA uptake are hindered. **(D)** When the negative regulator *npr1* is deleted, the inhibition on Art1’s function in endocytosis is removed. Under this condition, binding of Art1/Rsp5 to amino acid permeases is increased and Art1/Rsp5 are continuously recruited to the plasma membrane, which, in turn, stimulates amino acid permeases and plasmid DNA invagination.

Here, we investigated the roles of Art1, Art3, Bul1 and the ubiquitin ligase Rsp5 in facilitating the increase in transformation by targeted gene deletion and a complementation. The transformation efficiencies of *Δart1, Δbul1*, and *Δart3* cells were compared to that of wild type S. *cerevisiae* (Figure 3C). The median transformation efficiencies of *Δart1, Δart3* and *Δbul1* without amino acid addition was not significantly higher than those of wild-type yeast without amino acid addition (Tukey’s multiple comparisons test, p=0.5640 (wild-type vs *Δart1*), p=0.9195 (wild-type vs *Δart3*), p=0.9908 (wild-type vs *Δbul1*)). When amino acids were added to the *Δart3* and *Δbul1* strains, transformation efficiencies were substantially higher for both *Δart3* and *Δbul1* (up to about 20-fold; Tukey’s multiple comparisons test, p<0.0001). In contrast, as seen before, there was no boosting effect in the *Δart1* mutant (Tukey’s multiple comparisons test, p=0.9367). We deduce from this that TORC1-Npr1-Art1 signalling is specifically required for the boost in transformation efficiency induced by addition of amino acids to the media.

To test this hypothesis further, and to elucidate the role of the Rsp5, we carried out complementation of these mutants by 1) by the wild-type *art1* gene (pRS426-Ldb19) and a mutant *art1* gene from which the Rsp5-binding domain is deleted (pRS426-Ldb19PPxY-less) (Figure 2D). We found that *Δart1* was effectively rescued by the wild type gene: addition of amino acids significantly increased (over 24-fold; Tukey’s multiple comparisons test, p<0.0001). Conversely, there was no significant difference when *Δart1* was complemented by the gene lacking the Rsp5-binding domain (Tukey’s multiple comparisons test, p=0.6103). In summary, we found that the enhancement of transformation in response to addition of amino acids is mediated by TORC1-Npr1-Art1/Rsp5 (coloured) rather than other TORC1-Npr1-arrestins/Rsp5 routes (grey) (Figure 3D). Moreover, Rsp5 binding to Art1 is essential for this, which indicates that Rsp5-mediated mono-ubiquitination of plasma membrane cargo followed by ubiquitin-mediated endocytosis are necessary for enhanced yeast transformation.

**Figure 3.**
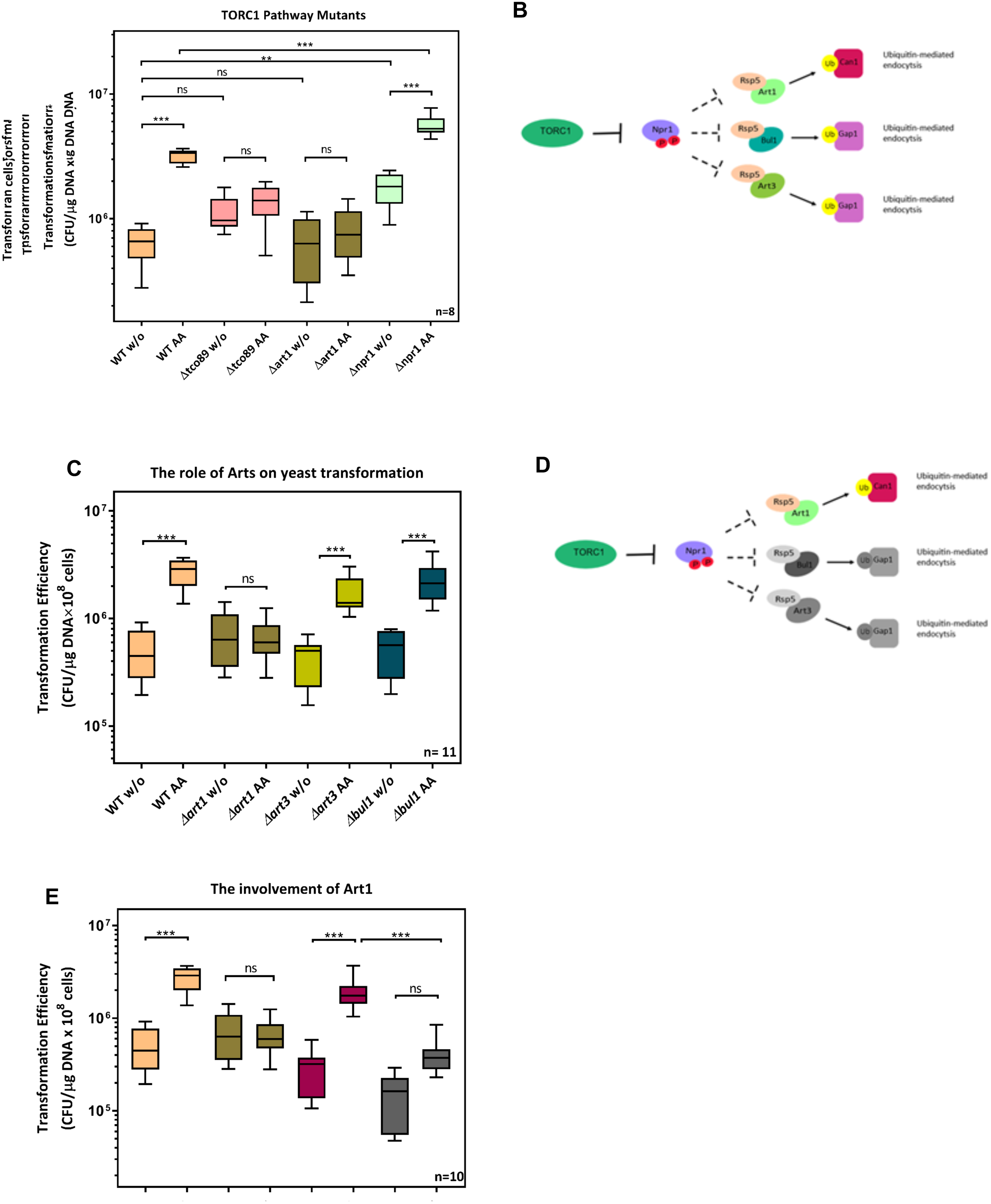
The TORC1 signalling pathway is involved in yeast transformation. **(A)** Three genes involved in TORC-1 signally were deleted from *S. cerevisiae* MaV203: the three strains generated were *Δtco89, Δart1*, and *Δnpr1*. Wild-type yeast (MaV203) and the mutant strains were transformed either with or without amino acid addition. The corresponding efficiencies were assessed by Tukey’s multiple comparisons test. The boosting effect was absent when either *tco89* or *art1* was deleted while transformation was enhanced when *npr1* was deleted. Results are from eight independent biological replicates. **(B)** The cartoon shows the pathways from TORC-1 to Npr1’s downstream targets, Art1, Art3, and Bul1. When TORC1 is active, the Npr1 kinase is inhibited, which allows Art1/Rsp5 binding to amino acid permease Can1, followed by Can1 invagination. When npr1 is inhibited, it also allows Art3/Rsp5 and Bul1/Rsp5 acting in ubiquitin-dependent cargo-selection of the general amino acid permease Gap1. **(C)** The requirement of different arrestins for transformability was tested. Three MaV203 mutants were generated, namely, *Δart1, Δart3, Δbul1*, and these mutants were transformed either with or without amino acid addition, followed by three-day culturing on 1x YPAD drop-out leucine and tryptophan. The corresponding transformation efficiencies were assessed by Tukey’s multiple comparisons test. Boosting was abolished only when Art1 was deleted suggesting that the effect is mediated by TORC1-Npr1-Art1 signalling route. Results are from eleven independent biological replicates. **(D)** The boosting effect is mediated by TORC1-Npr1-Art1/Rsp5 signalling (in colour) while both TORC1-Npr1-Art3/Rsp5 signalling and TORC1-Npr1-Bul1/Rsp5 signalling are not involved in boosting (in grey colour). **(E)** The Art1-PPxY-motif is required for effective plasmid DNA uptake. MaV203, *Δart1, Δart1* carrying pRS426-Ldb19 (*art1+*), and *Δart1* carrying pRS426-Ldb19PPxY-less (the PPxY motif required for Rsp5 binding was deleted from Art1) were transformed either with or without amino acid addition. After the transformation, the cells were cultured on 1x YPAD selection plates for three-days, followed by assessing the efficiencies by Tukey’s multiple comparisons test. The results demonstrated that complementation of Art1 in *Δart1* restored boosting. However, the effect was not observed when Art1 was expressed without the Rsp5 binding ability. Results are from ten independent biological replicates.

### Seg1 is required for high-efficiency yeast transformation

The integrity of eisosomes is known to affect the efficacy of endocytosis (Murphy, Boxberger et al. 2011). Here, we investigated the effect of MCC/eisosome formation on yeast transformation by deletion of *seg1* (known to impair the formation of eisosomes (Moreira, Schuck et al. 2012) and by deletion of *ypk1* (a kinase involved in eisosome formation (Luo, Gruhler et al. 2008)) (Figure 4). We found that removing *seg1* or *ypk1* resulted in no amino-acid induced increase in transformation efficiency (*Δseg1* without amino acid addition vs *Δseg 1* with amino acid addition; p=0.9873; *Δypk1* without amino acid addition vs *Δypk1* with amino acid addition, p=0.9976). It is noteworthy that although the boosting effect on both *Δseg1* or *Δypk1* disappeared, there were evident differences in the basal transformation efficiencies in the absence of added amino acids to the media (wild-type vs *Δseg1*: p=0.0249; wild-type vs *Δypk1:* p=0.0118).

**Figure 4.**
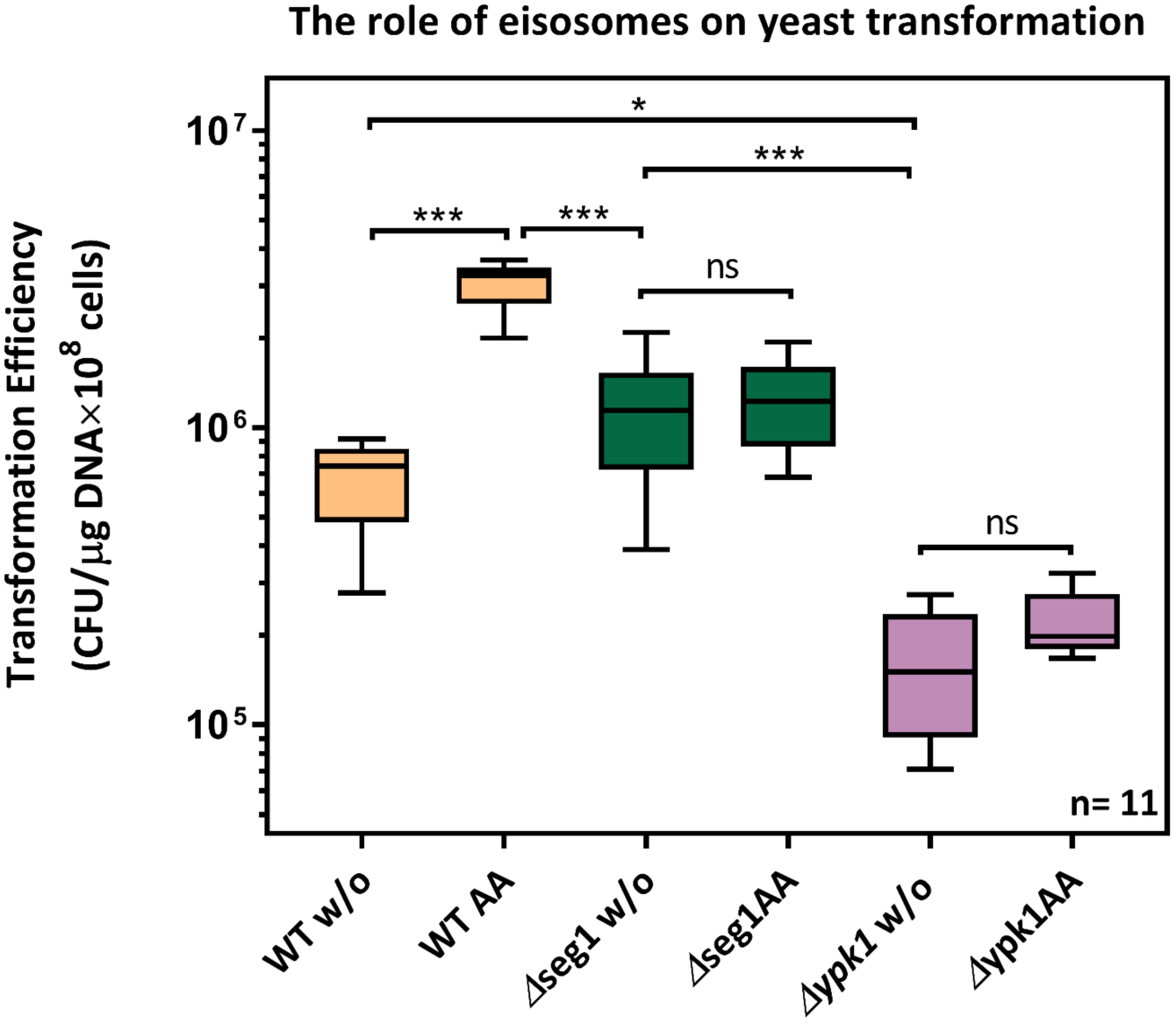
Eisosomes are required for increased transformation efficiency in yeast. *Δseg1* and *Δypk1* strains generated by deletion of the respective genes were transformed either with or without amino acid addition to the media. The corresponding transformation efficiencies were analysed by Tukey’s multiple comparisons test. The results showed that the boosting effect is abolished in both *Δseg1* and *Δypk*1, suggesting that functional eisosome formation and signalling are required for highly efficient yeast transformation. Results are representative of 11 independent biological replicates.

## Discussion

Yeast transformation has been described for almost 40 years (Hinnen, Hicks et al. 1978) and is a cornerstone of many fundamental methods in genetics, cell biology as well as practical biotechnological applications. It is therefore surprising there are only few mechanistic explanations of the processes underpinning this key technique although several have been proposed (Beggs 1978). One model suggests that foreign DNA is engulfed via endocytic membrane invagination; this is supported by the observation that several low transformability phenotypes are caused by mutation of genes involved in endocytosis (Kawai, Pham et al. 2004). Here, we tested the extent to which targeted deletions of single endocytic genes affect competence, and we observed how the changes to nutrients in the growth and the transformation media affected transformation efficiency. We found that adding amino acids boosted competence in all four strains of *S. cerevisiae* tested, demonstrating that this phenomenon is likely to be generally applicable in budding yeast. We propose a model to summarise the processes described here (Figure 5).

**Figure 5.**
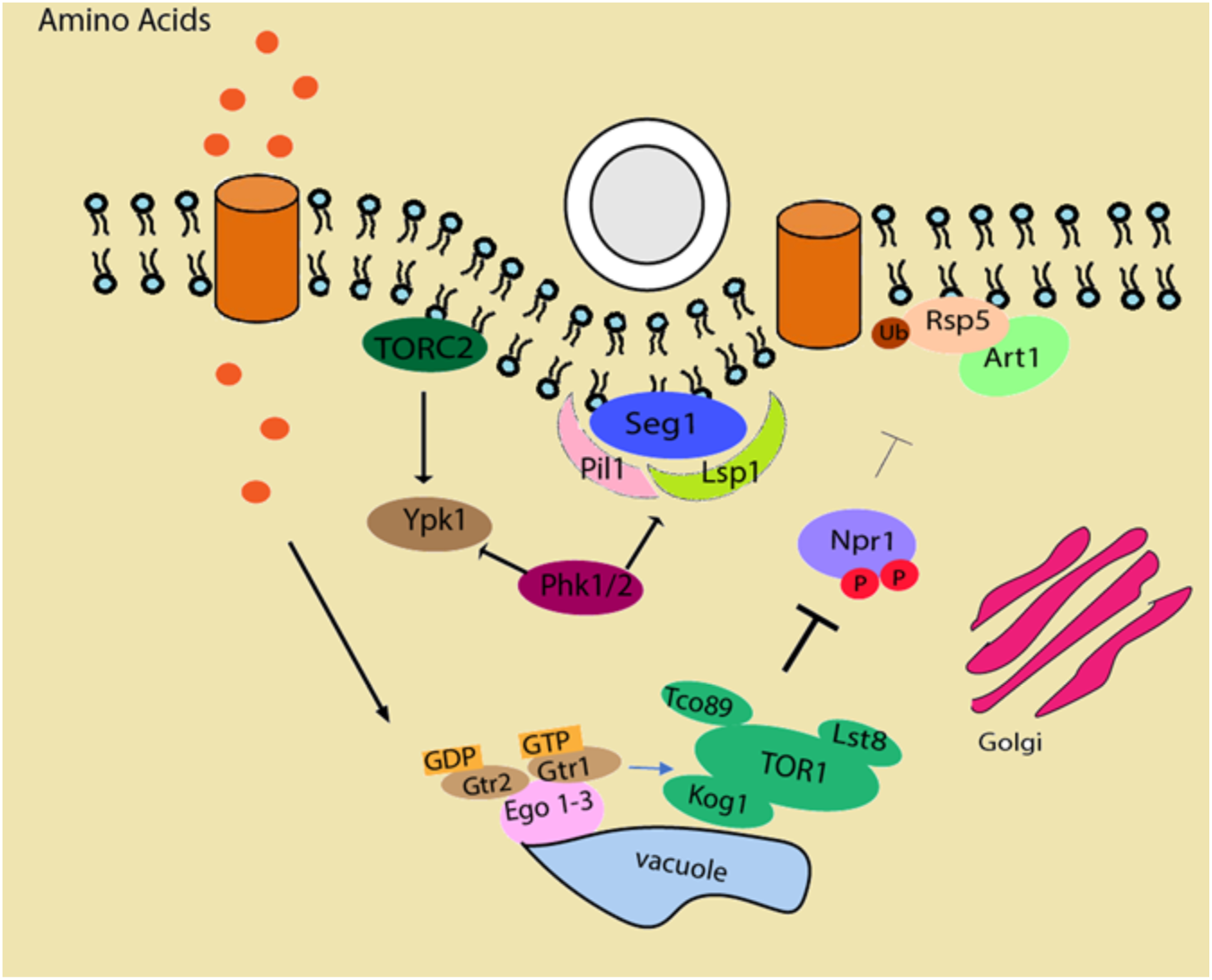
Proposed model of TORC1-regulated DNA uptake in yeast and the involvement of eisosomes. Amino acids are transported into the cell cytoplasm via amino acid permeases, where they stimulate TORC1. Activated TORC1 inhibits the Npr1 which, in turn, facilitates Art1/Rsp5-dependent cargo selection and subsequent endocytosis with simultaneous plasmid DNA uptake. Seg1, Pil1 and Lsp1 are the main proteins involved in eisosome formation and stability. Phk1/2 phosphorylates ypk1 and seg1 this initiates the deposition of Pil1/Lsp1 where eisosomes form. Following this, phosphorylation of Pil1 stabilises the formation of eisosomes, which, in turn, enhances the rate of plasmid DNA internalisation.

### TORC1 and other pathways impinge on transformation efficiency

The *tco89* gene encodes a subunit of TORC1. We observed that in *Δtco89* cells there was never any boosting effect in response to nutrient stimuli. However, deletion of *tco89* did not lead to changes in basal competence, but only affected the boost induced by amino acids in the media. Therefore, whileTORC1 is necessary for the regulation of the amino acid-induced effect, there are also other pathways underpinning DNA uptake. Plasma membrane permeases such as Can1 are under the control of TORC1, but are also internalized in response to their substrate, independently of TORC1 signalling (Opekarova, Caspari et al. 1998, Ghaddar, Merhi et al. 2014). Our results imply that, in contrast to TORC1-regulated internalization, the influx-stimulated internalization of PM permeases does not contribute to efficient DNA uptake or subsequent delivery to the nucleus.

In contrast, the basal competence of *Δnpr1* cells was higher than wild type. This was attributed to abolition of the of α-arrestins’ inhibition of these endocytic processes (MacGurn, Hsu et al. 2011). The boost of transformation observed in *Δnpr1* cells treated with amino acids reveals there are other regulators of arrestins under the control of TORC1, in addition to Npr1.

### Efficient DNA delivery requires functional Art1

We discovered that a functional Art1 was indispensable for high transformability in response to amino acids. In experiments where we complemented the *art1* deletion, we also observed that the Rsp5-binding domain of Art1 was required for the boosting effect. Because of this, we propose that ubiquitin-mediated cargo sorting is involved in efficient DNA uptake *in vivo*.

While Art1 was needed for high transformability in response to amino acids, deletion of other arrestins that act in a similar manner (Bul1, Art3) had no effect. This is important as both Bul1 and Art3 are under the control of Npr1 and are involved in endocytosis (O’Donnell, Apffel et al. 2010, O’Donnell 2012). Thus, the TORC1 – Npr1-Art1 pathway is specifically responsible for the effect on competence investigated here. This might be explained by the observation that the three arrestins we tested have different targets localised to different domains of the plasmamembrane. For example, Art1 is involved in Can1 endocytosis, whereas Bul1 and Art3 function in Gap1 down-regulation and recycling (Helliwell, Losko et al. 2001, Lin, MacGurn et al. 2008, O’Donnell, Apffel et al. 2010). In the plasma membrane, whilst Gap1 is uniformly distributed (Lauwers, Grossmann et al. 2007), there are specific compartments containing Can1 (MCC) (Nikko and Pelham 2009): therefore, these results hinted that MCC may be important domain in transformation competence.

### Involvement of eisosomes in yeast transformation

Eisosomes are cytosolic multi-protein complexes that form 50–300 nm deep invaginations of the plasma membrane associated with the MCC domain (Stradalova, Stahlschmidt et al. 2009). The membrane composition of the MCC differs from other yeast membrane compartments because it contains more ergosterol (Grossmann, Opekarová et al. 2007). Eisosomes act as a hub for various signalling pathways and may play a role in endocytosis (Walther, Brickner et al. 2006, Fröhlich, Moreira et al. 2009, Douglas and Konopka 2014). Research on endocytic activity associated to eisosomes has led to several intensely debated results (Grossmann, Malinsky et al. 2008, Vangelatos, Roumelioti et al. 2010, Brach, Specht et al. 2011, Murphy, Boxberger et al. 2011, Seger, Rischatsch et al. 2011, Athanasopoulos, Boleti et al. 2013). Eisosomes require *seg1* for stability (Moreira, Schuck et al. 2012).

Our finding that mutant strains lacking *seg1* do not show high transformability in response to an amino acid stimulus (Figure 2E), is indirect support for claims that eisosomes either mark sites of endocytosis or positively regulate endocytosis (Walther, Brickner et al. 2006, Murphy, Boxberger et al. 2011). Importantly, although the basal competence of *Δseg1* cells is similar to wild type, the role of eisosomes in transformation is unlikely to be restricted to the boosting effect, because a subset of eisosomes still forms in *Δseg1* cells (Moreira, Schuck et al. 2012).

One of the functions of *YPK1* kinase is to control eisosome formation (Luo, Gruhler et al. 2008). Deletion of *YPK1* led to the lowest transformation efficiencies out of all mutant strains we tested in this study. However, further work is needed to clarify the exact role of Ypk1 in DNA uptake because: Ypk1 also regulates at least one α-arrestin (Alvaro, Aindow et al. 2016), it impinges on actin dynamics (Niles and Powers 2014), and it is involved in the heat stress response (Sun, Miao et al. 2012). The hypotheses that eisosomes mark sites of endocytosis and that transformation is facilitated by endocytosis have one point in common: in both cases these types of endocytosis differ from well-studied endocytic pathways, such as clathrin-mediated endocytosis that originates at actin patches (Kawai, Pham et al. 2004, Ziółkowska, Christiano et al. 2012).

Alternative endocytic pathways in yeast are not as well-studied as in mammalian cells. New insights emerged in recent years, for example the α-arrestins Art1 and Bul1 can lead to endocytic downregulation of transmembrane transporters in a clathrin- and ubiquitin-independent manner, relying on Rho1 (Prosser, Drivas et al. 2011, Prosser, Pannunzio et al. 2015). Indeed, as stated above, we found that the Rsp5-binding domain in Art1 was required to observe high transformation efficiency. This implies that ubiquitination of cargo proteins does at least partially contribute to subsequent DNA uptake. Whether endocytic DNA uptake relies on clathrin-coated vesicles and ubiquitination as a cargo signal *per se*, remains to be seen. Nevertheless, we propose that an eisosome-mediated pathway is the main route for efficient DNA delivery into the yeast cell.

## Outlook

A complete mechanistic description of the genetic requirements and endocytotic mechanism for nucleic acid uptake is immensely important not just for understanding yeast transformation but also for further progress in various fields like gene therapy in humans, understanding RNA trafficking and improving RNA interference technologies.

It will be exciting to see whether the concept of achieving high transformation efficiencies in yeast by stimulation of TORC1 can be applied to the mammalian mTORC system as well. By highlighting the importance of the metabolic state of the cell, this study opens up new practical possibilities for the improvement of transformation efficiencies, by fine-tuning the nutrient composition in the transformation reagent.

## MATERIALS AND METHODS

### *S. cerevisiae* strains, plasmids, reagents and equipment

This study includes an evaluation of transformation efficiencies of four *S. cerevisiae* strains, under different nutrient conditions. Namely, the yeast strains MaV203 (from ProQuest™ Two-Hybrid system (PQ10001–01, Thermo Fisher Scientific), W303–1A, BY-4743, and THY.AP4 (kindly provided by Bjorn Sabelleck, RWTH Aachen University). The MaV203 strain was used to generate seven mutant strains. These included *Δart1, Δart3, Δbul1, Δnpr1, Δseg1, Δtco89*, and *Δypk1*. Four plasmids, namely, pDEST22 (PQ1000101, Thermo Fisher Scientific), pDEST32-TaRNR8-p12L (generated by Dr Sheng-Chun Yu), pRS426-Ldb19, and pRS426-Ldb19^PPxY-less^ the last two plasmids kindly provided by Allyson F. O’Donnell, Duquesne University, PA USA were used in this study. An AccuTherm™ Microtube Shaking Incubator (I-4002-HCS, Labnet International, *Inc*.) was used for the heat-shock process in yeast transformation. We used the following reagents in this study: Yeast extract (Y1625–250G, Sigma-Aldrich), Peptone (P5905–1KG, Sigma-Aldrich), Adenine hemisulfate salt (A3159–100G, Sigma-Aldrich), D-(+)-Glucose (G7021–1KG, Sigma-Aldrich), yeast nitrogen base without amino acids (Y0626–250G, Sigma-Aldrich), yeast synthetic drop-out medium supplements (Y2001–20G, Sigma Aldrich), L-histidine monohydrochloride monohydrate (53370–100G, Sigma-Aldrich), L-tryptophan (T8941–25G, Sigma-Aldrich), uracil (U1128–25G, Sigma-Aldrich), D-sorbitol (S3889–1KG, Sigma-Aldrich), Poly(ethylene glycol) BioUltra, 1000 (PEG1000) (81188–250G, Sigma-Aldrich), LiAc (6108–17–4, Alfa Aesar), Deoxyribonucleic acid sodium salt from salmon testes (ss-DNA) (D1626–5G, Sigma-Aldrich), Bicine (B3876–100G, Sigma-Aldrich), ethylene glycol(324558–100ML, Sigma-Aldrich), dimethyl sulfoxide (DMSO) (D2650–5 × 5ML, Sigma-Aldrich), Water Molecular Biology Reagent (W4502, Sigma-Aldrich), Ultra-Pure™ Agarose (16500500, Thermo Fisher Scientific), SYBR^®^ Safe DNA Gel Stain (S33102, ThermoFischer Scientific), GeneRuler 1kb Plus DNA ladder (SM0311, ThermoFischer Scientific), Zymolyase from Easy Yeast Plasmid Isolation Kit (630467, Clontech), GoTaq^®^ G2 DNA Polymerase and 5X Colorless GoTaq^®^ Reaction Buffer (M7841, Promega) and dNTP mix (R0191, ThermoFischer Scientific).

### *S. cerevisiae* transformation

*S. cerevisiae* transformations were performed using the SuccessAA protocol (Yu, Dawson et al. 2016), an adaptation of the LiAc/SS carrier DNA/PEG method (Gietz 2015) with the addition of amino acids in the transformation mix. The concentration of amino acids used was 1.25X the concentration of amino acids found in Synthetic Complete (SC) medium. Briefly, 0.25 µg endotoxin-free pDEST32-TaRNR8-p12L plasmid (13.8kb) was added into 50 µl MaV203 competent cells, followed by adding 500 µl transformation mix solution, containing 36% (w/v), PEG 1000, 0.1 M LiAc, 0.2 mg/ml ss-DNA, 0.2 M Bicine-NaOH (pH=8.35), and 1.25x amino acid mix solution. The plasmid DNA was mixed to the competent cells in the transformation mix solution, and the yeast cells were then heat-shocked in the microtube shaking Incubator at 37°C for 30 minutes. The transformation mixtures were shaken at the start and after 15 minutes at 1500 rpm for 5 seconds; at the end of the incubation, the samples were shaken for 30 seconds. After the heat shock, 50 µl transformation mixtures containing wild type MaV203 yeast cells or different mutated MaV203 yeast strains were plated on suitable synthetic “drop-out” plates, and cultured for three days at 30°C. The numbers of colony forming units (CFU) were counted, and transformation efficiencies (E) were calculated with the following formula:

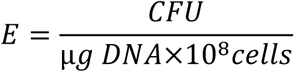

### *S. cerevisiae* mutant strains generation and yeast colony PCR

The current study generated seven yeast mutants to investigate the potential molecular mechanisms underlying yeast transformation. Targeted gene deletion mutagenesis (gene “knock-out”) mediated by homologous recombination reaction was used to mutate the following genes in MaV203: *art1, art3, bul1, npr1, seg1, tco89*, and *ypk1*. Mutagenesis primers were designed so that the gene of interest would be replaced by *TRP1*, which served as an auxotrophic selection marker carried by the pDEST22 plasmid. Primer sequences are shown in Supplementary Table 1. All the sequences of the forward primers were 74 bases whereas the first 50 bases were identical to the first 50 bases of the target gene, followed by the reverse and complementary 24-base sequence (6431bp to 6454bp on pDEST22) adjacent to the ARS/CEN locus in the pDEST22 plasmid. Similarly, the sequences of the reverse primers were 74 bases whereas the first 50 bases were identical to the last 50 bases of the target gene, followed by the reverse and complementary 24-base sequence (5143bp to 5166bp) which is adjacent to the f1 origin in the pDEST22 plasmid. The plasmid pDEST22, carrying *TRP1*, was used as a PCR template. The PCR thermal cycled we used was: initial denaturation at 95°C for 3 minutes, followed by 40 cycles of denaturation at 95°C for 30 seconds, annealing at 45°C for 30 seconds, and extension at 72°C for 2 minutes and then the final extension at 72°C for 7 minutes. The PCR products were examined by gel electrophoresis. Once the PCR products exactly matched the predicted size, the PCR product was purified using QIAquick PCR Purification Kit. The gene specific PCR products were used to transform S. *cerevisiae* MaV203 as as described above. A 100 µl aliquot of the transformation mixture was plated on synthetic complete “drop out” tryptophan plates, followed by culturing the plates at 30°C for three days. Potentially mutated MaV203 yeast colonies were analysed using a modified version of yeast colony PCR protocol published in *Molecular Cloning: A Laboratory Manual* (Green 2012). In brief, at least ten colonies on each plate were randomly selected and approximately 1/10^th^ of each colony was carefully transferred to each sterile PCR tube, containing 5 µl zymolyase solution (from Easy Yeast Plasmid Isolation Kit). The PCR tubes with yeast-zymolyase mix were then incubated for 30 minutes at 37°C, followed by incubating for 10 minutes at 95°C to inactivate zymolyase. After zymolyase inactivation, the yeast-zymolyase mix was diluted by addition of 95 µl molecular biology grade endotoxin-free water and then yeast colony PCR was performed. The yeast colony PCR program was as follows: initial denaturation at 95°C for 5 minutes, followed by 40 cycles of denaturation at 95°C for 1 minute, annealing at 53°C for 1 minute, and extension at 72°C for 2 minutes and then the final extension at 72°C for 7 minutes. When the colony PCR finished, 5 µl PCR reactions were analysed on a 1.5% (w/v) agarose/TBE gel for 45 minutes at 10V/cm. Successful transformants were identified based on the predicted PCR product length, for that primers of the adjacent down- and upstream region of the target gene where designed. Reverse primers binding to *TRP1* were used in a separate PCR reaction for an additional verification.

### Evaluation of transformation efficiencies

Once the mutants were confirmed by yeast colony PCR, the mutants were cultured on 1x YPAD “drop-out Tryptophan” plates for three days, followed by growth in the same 1xYPAD “drop-out” medium overnight. Frozen yeast mutant competent cells were then prepared, transformed, and cultured on 1xYPAD “drop-out leucine and tryptophan” plates using the SuccessAA protocol (Yu, Dawson et al. 2016). Mutant transformation efficiencies were measured and compared to the efficiency of wild type MaV203 yeast cells.

### Statistical Analysis

The collected data was not always normally distributed so non-parametric tests were used, where appropriate. In the evaluation of the transformation efficiency of different budding yeast strains, there was no inter-group comparison, only two groups were compared at a time (same yeast strain, with the addition of amino acids or without). Thus, the Mann-Whitney test were used to assess statistical significance. For assessment of APIU’s effect on transformation efficiency, there were three groups to be compared so a Kruskal Wallis test, followed by Dunn’s post-hoc test was used. In the rest of the experiments, there was an interaction between the two factors tested, the addition of amino acids and different mutant yeast strains. For this reason, two-way ANOVA was used to assess statistical significance, followed by Tukey’s multiple comparisons test. Statistical analysis was performed in GraphPad Prism 7.03.

## Contributions

S.-C.Y. conceived the original idea of this study. S.-C.Y., F.K. and N.S.P. designed and carried out the experiments, analysed the data and drafted the paper. P.D.S. supervised and advised the work and edited the manuscript.

## Acknowledgements

We thank Mark Isalan (Imperial College London) for invaluable advice on yeast mutagenesis Allyson O’Donnell (Duquesne University, PA USA) who kindly provided and the pRS426-Ldb19, and pRS426-Ldb19^PPxY-less^ plasmids.

---

**Supplementary Table S1:**
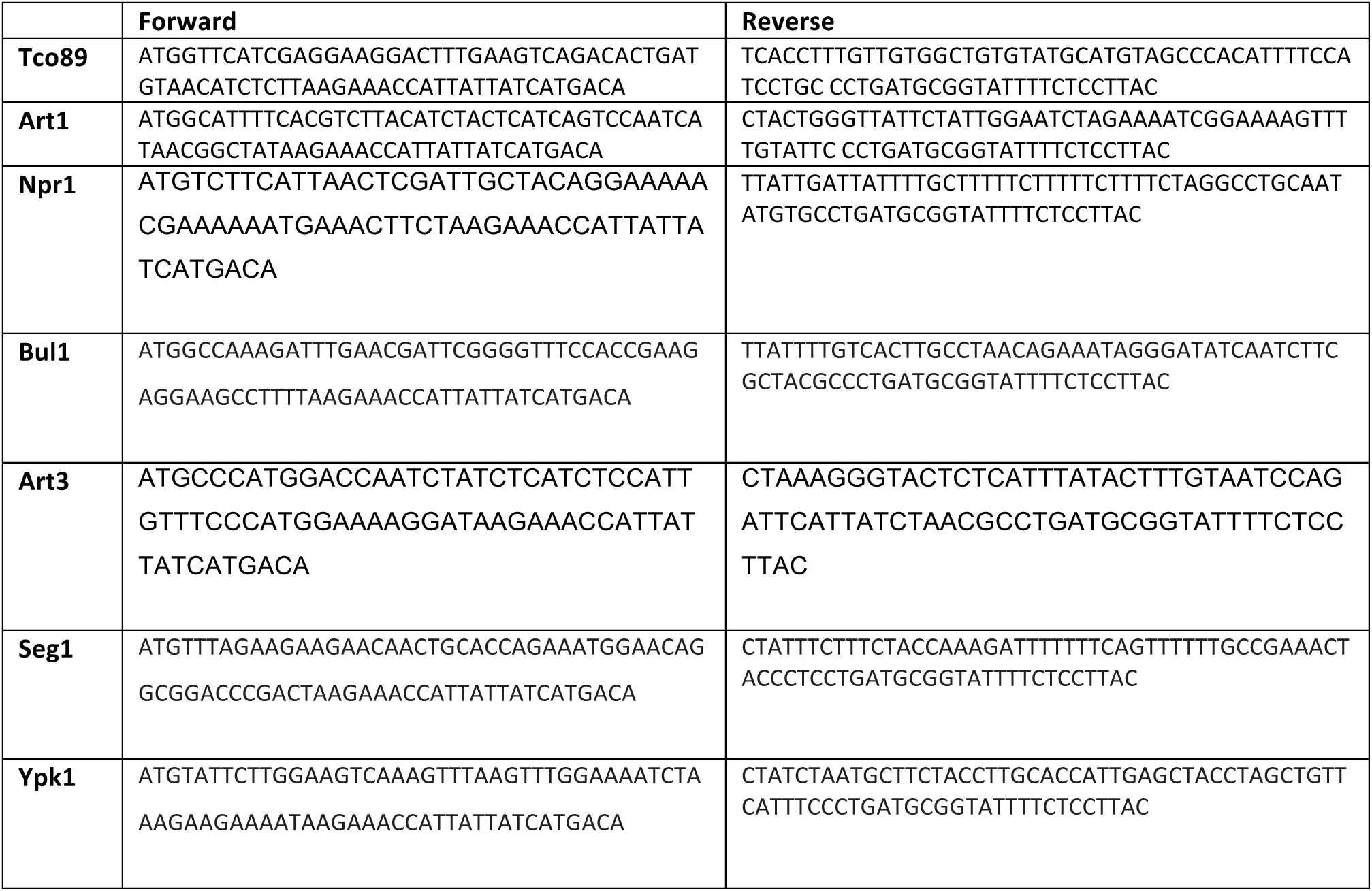
Mutagenesis primers sequence

**Supplementary Table S2:**
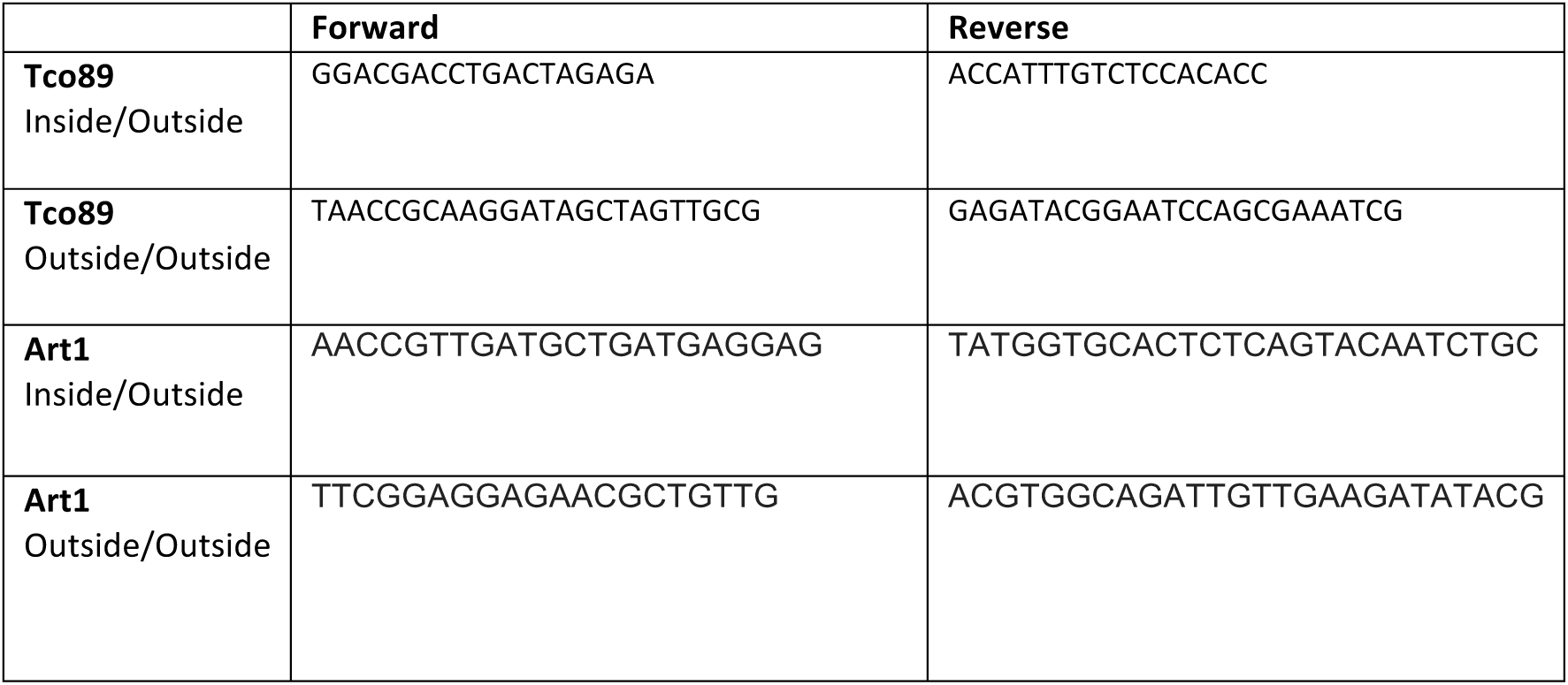
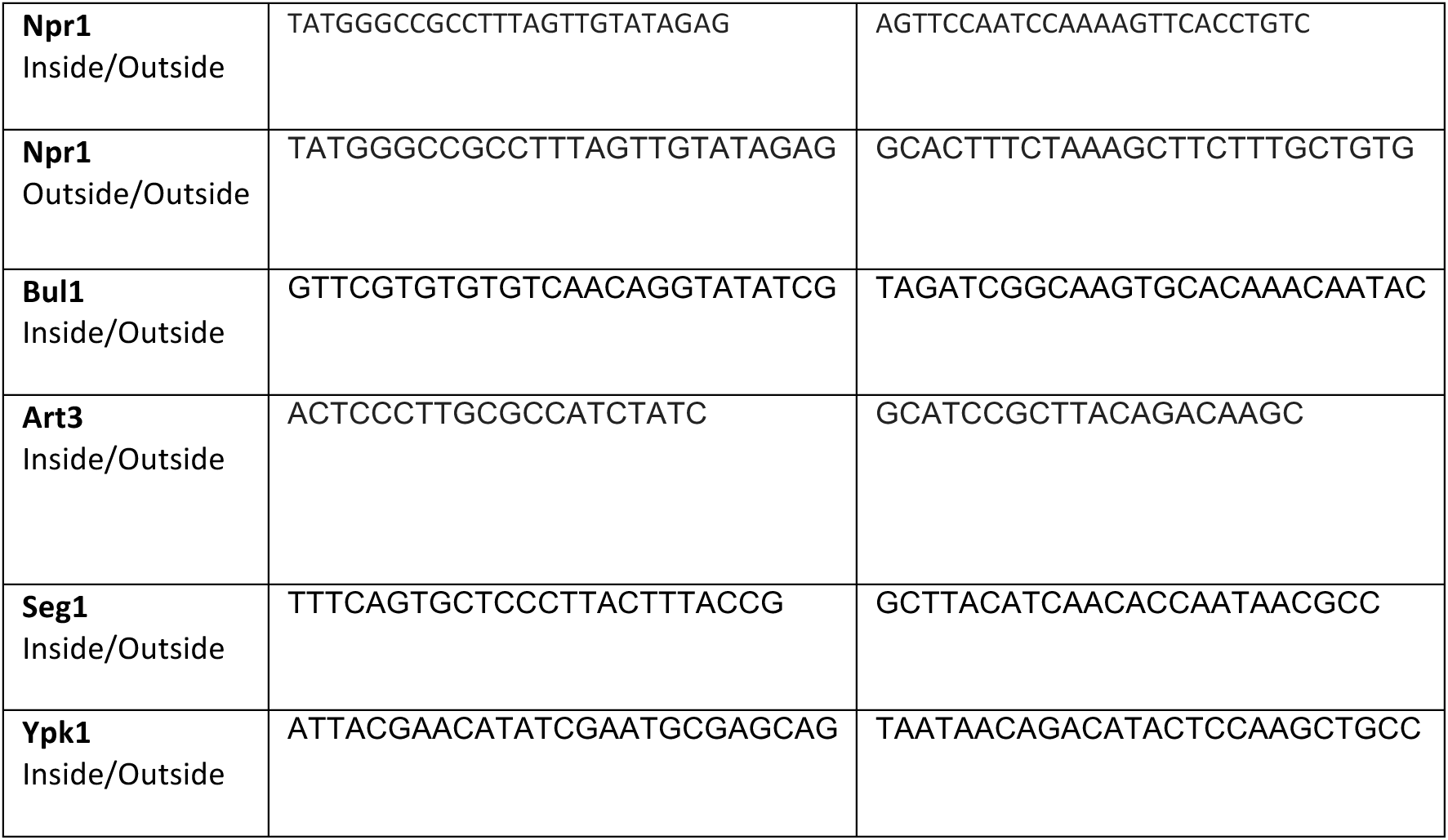
Diagnostic primers sequence

